# Conformable and robust force sensors to enable precision joint replacement surgery

**DOI:** 10.1101/2021.08.19.456934

**Authors:** Liam Ives, Alizée Pace, Fabian Bor, Qingshen Jing, Tom Wade, Jehangir Cama, Vikas Khanduja, Sohini Kar-Narayan

## Abstract

Balancing forces within weight-bearing joints such as the hip during joint replacement surgeries is essential for implant longevity. Minimising implant failure is vital to improve patient wellbeing and alleviate pressure on healthcare systems. With improvements in surgery, hip replacement patients are now often younger and more active than in previous generations, and their implants correspondingly need to survive higher stresses. However, force balancing currently depends entirely on surgical skill: no sensors can provide quantitative force feedback within the hip joint’s small, complex geometry. Here, we solve this unmet clinical need by presenting a thin and conformable microfluidic force sensor, which is compatible with the standard surgical process. We optimised the design using finite element modelling, then incorporated and calibrated our sensor in a model hip implant. Using a bespoke testing rig, we demonstrated high sensitivity at typical forces experienced during hip replacements. We anticipate that these sensors will aid implant positioning, increasing the lifetime of hip replacements, and represent a powerful new surgical tool for a range of orthopaedic procedures where force balancing is crucial.

---

Over 1.9 million total hip replacements (THRs) and 1.5 million total knee replacements (TKRs) were performed in the United Kingdom between April 2003 and December 2019^1^. This rate is ever increasing, with over 280,000 THRs and 320,000 TKRs performed between 2017 and 2019. In the United States, the demands for hip and knee replacements are predicted to rise to 1,429,000 and 3,416,000 annual cases respectively by 2040^2^. This is due to both an ageing global population and joint replacement patients being younger and more physically active than a few decades ago^2,3^. The greater the number of joint replacements, the greater the number of resultant revision procedures required when the primary implant fails. Overall, 10% of primary THRs between April 2003 and December 2019 have required revision^1^, but the failure rate increases to 42% 25 years after primary hip surgery^4^. Revision surgery is expensive and technically challenging, so there is considerable interest in improving surgical procedures to maximise the implant longevity^5–11^. Implant survival or longevity is based on several factors, but implant positioning plays a fundamental role. Poor implant positioning has been correlated with a higher rate of dislocation, prosthetic impingement, leg length inequality, implant loosening, accelerated wear, poor functional outcomes, and finally increased revision rates^12–14^. These comprise about 40% of the causes of revision surgery^15^. The primary THR dislocation rate can be as high as 7%, whereas this rate increases to 28% for revision THRs^1^. Restoration of normal hip anatomy at the THR provides for better clinical function and abductor strength as well as reduced wear of the polyethylene implant component which is prone to failure^16–20^.

The hip is a ball and socket joint, and the biomechanical goals of a total hip replacement are to achieve the correct centre of rotation of the femoral head, and accurate leg length, offset and positioning of the femoral and acetabular components allowing hip stability. During a total hip replacement, the surgeon has access to the both the acetabulum, i.e. the socket – cup shaped part of the pelvic bone, and the femoral head, i.e. the top of the thigh bone, the femur. After removing the damaged cartilage and bone, the acetabular part of the implant, a metal or polyethylene cup, is inserted into the acetabulum. For a metal cup, an ultra-high molecular weight polyethylene (UHMWPE) cup is inserted, to provide a low-friction articulating surface to contact the implant’s femoral head. A metallic stem is inserted into the hollowed-out femur. A trial femoral head is attached to the top of the stem, and together the femoral and acetabular components are connected. The cup, head and stem components come in a range of sizes, so the implant size can be tailored to the patient’s joint. The surgeon then assesses the position, stability, leg-length and soft tissue balance of the trial components by checking for combined anteversion, moving the hip through a full range of movement but especially into flexion, internal rotation, extension, external rotation and telescoping the joint. Implant repositioning and resizing are performed if required and the process of trialling is repeated. Once the trial implants are deemed to have been correctly positioned with the correct soft tissue tensioning, the trial implants are exchanged with the final implants.

It is during this trial stage that surgeons lack objective force data to aid precise positioning and balance. Quantitative force feedback could provide information on whether the cup and stem are the correct distance apart (the correct offset) which leads to accurate soft tissue balance and whether this offers maximum stability to the joint. Furthermore, dynamic assessment of the joint during the trialling process with accurate force feedback could also indicate the uniformity of force distribution across the joint. This will also inform on the peak forces in cases of prosthetic impingement. This force balancing is crucial to restoring the anatomy of the joint, to allow optimal function in terms of range of movement and pain reduction; it is also necessary to reduce the chances of dislocation, prosthetic impingement, and wear over time.

In the knee, the lack of quantitative force feedback during implant positioning to inform the surgeon on balancing and implant size choice has recently been addressed to an extent with the single-use VERASENSE sensor (Orthosensor Inc.)^23^. This technology uses flat, rigid piezoelectric elements to provide force and positioning information during a TKR^24^. However, this cannot be readily adapted for the hip joint as the hip joint is curved and has much less space than the knee. Any solution for the hip (or other ball-and-socket joints) requires components that are thin, flexible, but still capable of bearing heavy loads.

This lack of objective measurement of force distribution during the critical part of a hip replacement is a major unmet need, and addressing it has the potential to significantly reduce the number of hip revisions due to wear, dislocation, inadequate soft tissue balancing or prosthetic impingement.

To meet this need, we have developed a thin, flexible, biocompatible microfluidic force sensor that can be incorporated into the implant geometry during the trialling stage of performing a THR. We have also designed and fabricated a biomimetic trial insert to host an array of sensors in the acetabular part of the total hip replacement, which we use to model a 3-dimensional representation of the forces in a cup as the surgeon inserts the femoral head and loads the hip at different angles. We designed the sensor to meet the requirements for force measurements in a THR, but the design can be rapidly customised to target a range of biomedical force sensing applications.

## Results

### Sensor Design and Operation

Fig. 1a is a schematic of the sensor, whose design is based on the group’s previous work^25^. The sensor consists of (1) a microfluidic chip (Fig. 1b) with an embedded microfluidic channel (20 mm x 0.75 mm x 0.3 mm) and fluid reservoir (2 mm x 2 mm x 0.3 mm), and (2) an electrode layer (Fig. 1c) comprising interdigitated silver electrodes on a flexible polyimide (PI, Kapton) substrate for mechanical support. The chip can be made using a silicone elastomer such as polydimethylsiloxane (PDMS) poured into a stereolithography (SLA) 3D-printed mold, or directly SLA 3D-printed by photo-curing an elastomeric resin (Flexible Resin, Formlabs). The channel is aligned above the electrodes and is open to air at the end opposite to the reservoir. The reservoir is the active sensing area. It has a square cross section, is filled with fluid by a syringe, and can contain internal columns for mechanical support. The fluid is a 2:1 volume mixture of glycerol and deionised (DI) water, to balance the low volatility and high dielectric constant^25^. The interdigitated electrodes are made by aerosol jet printing (AJP) silver nanoparticle ink onto Kapton and are protected by a printed PI layer. The chip and electrode layer are bonded using laser-cut double-sided tape. Fig. 1d shows a photograph of the sensor, which has dimensions of approximately 30 mm x 5 mm x 1 mm and can be freely flexed (Fig. 1e) and incorporated into the THR UHMWPE insert without significant impact on sensor performance.

**Figure 1.**
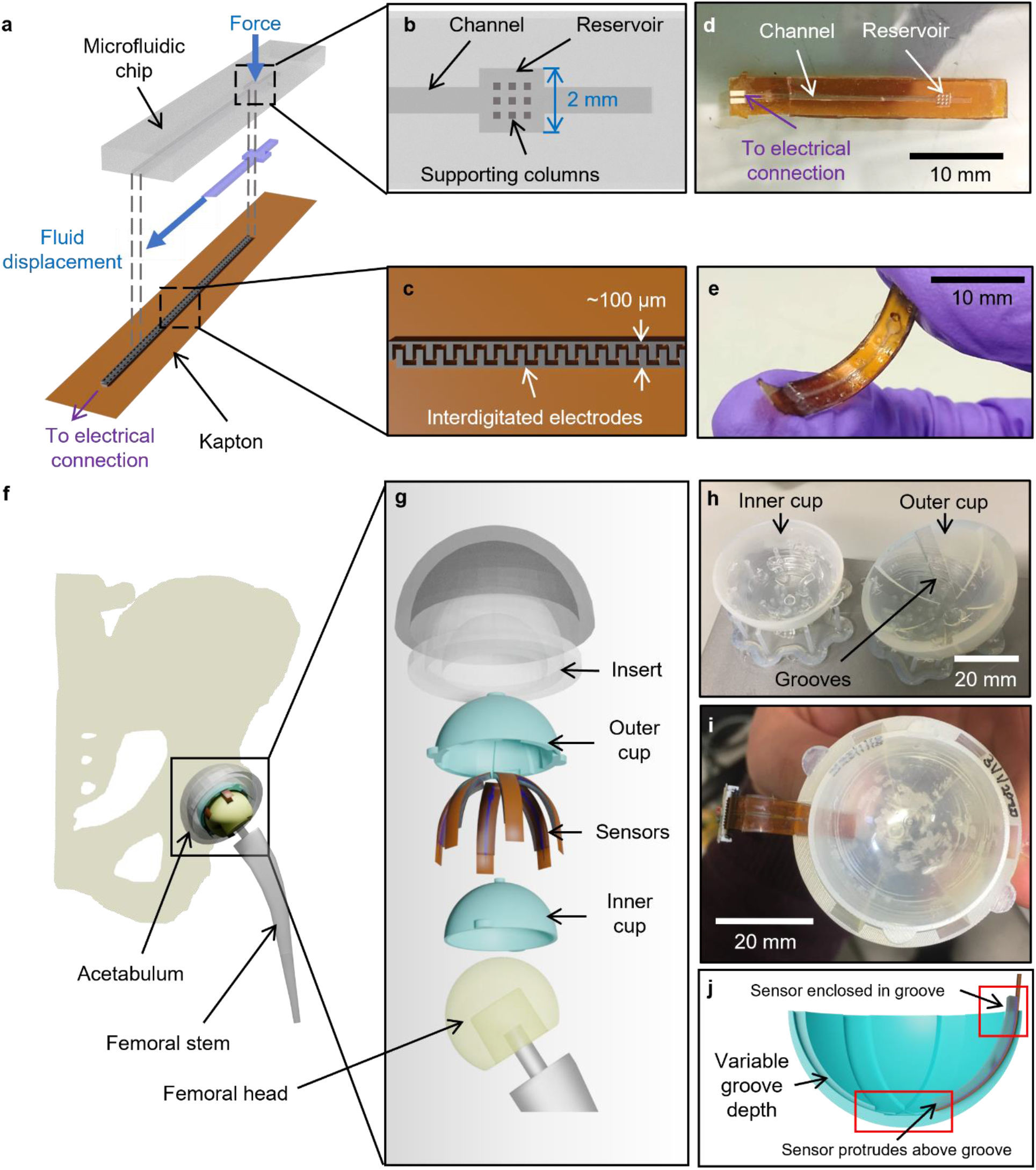
Incorporating functionalised microfluidic sensors into the THR implant for force measurements. **a** The sensor is made of a soft elastomeric microfluidic chip layer and a Kapton substrate with aerosol-jet printed interdigitated electrodes. **b** The microfluidic chip layer contains an embedded microchannel with a fluid reservoir and optional supporting columns. **c** Interdigitated electrodes are aerosol jet printed onto the Kapton substrate and consist of silver with an insulating polyimide coating. **d** Photograph of the sensor, highlighting the channel and reservoir. **e** Photograph of sensor, showing its flexibility. In this design, the reservoir has a round cross-section. **f** Standard geometry of a hip implant. The cup containing the sensors is incorporated into the polymer part of the implant’s acetabular cup. **g** Our additions to the implant, during the trial stage, consist of an outer cup, the sensors which lie in the grooves of the outer cup, and the inner cup to act as an articulating surface with the femoral head. **h** Photograph of the SLA 3D-printed inner and outer cup. **i** Photograph of a sensor located in a groove between the inner and outer cups. **j** Cross-section of the outer cup, showing how variable groove depth allows the channel to be shielded from force while the active sensing area (reservoir) is exposed.

The operating principle is as follows^25^: when a force is applied to the fluid reservoir, the reservoir deforms and displaces fluid along the channel. The displaced fluid overlaps with the interdigitated electrodes, increasing the capacitance. The capacitance is calculated using the equation *C* = *ε*_0_*ε*_*r*_*A*/*d*, where *ε*_*r*_ is the relative dielectric permittivity, *A* is the electrode area and *d* is the inter-electrode distance. The fluid determines the value of *ε*_*r*_. On releasing the force, the fluid returns to the reservoir.

The sensors are durable, with measurements reproduced over more than 2,000 loading cycles in a mechanical testing rig^25^. For each sensor, the force-capacitance relationship is calibrated by applying a known force and measuring the corresponding impedance using an impedance analyser (ISX-3, Sciospec), which is converted to a capacitance. For the present application, an array of sensors is embedded into the UHMWPE component.

Importantly, these sensors can be easily and rapidly customised to suit a range of applications by changing the reservoir and channel size, electrode size and spacing, chip material and device size. While this sensor has been developed for hip applications, we have fabricated sensors with alternative designs, including a sensor incorporating three sensing elements intended for the knee joint both for the TKR and unicompartmental applications.

### Incorporation into Implant

To test the sensor’s performance in the hip, a biomimetic trial insert (Fig. 1f) was designed using Creo Parametric (PTC) and produced by SLA 3D printing. The insert (Fig. 1g,h) consists of an ‘outer cup’, which contains grooves to fit sensors; and an ‘inner cup’, which has a smooth inner surface to act as an articulating surface with the femoral head, and pegs to secure it in the outer cup. It was decided to incorporate six sensors into the trial insert to provide a balance between providing many sensing regions and mechanical stability when loading the sensors. The sensors were incorporated into the grooves (Fig. 1i) such that the reservoirs were at 30° to the cup axis. In this study, the outer cup has an external diameter of 43 mm and an internal diameter of 41 mm, but this can be adapted to suit femoral heads of different sizes. Typically, the femoral head has a diameter between 28 and 40 mm^1^. The groove depth is deliberately reduced going from the outside of the cup towards the centre (Fig. 1j). This gives the effect of raising the reservoirs out of the grooves, such that an applied force is concentrated on the fluid reservoirs (the active sensing area) while shielding the channel.

Formlabs’ Durable Resin was used as the cup material – the resin has similar mechanical properties to the UHMWPE used in the acetabular insert of real hip implants (see Supporting Information Fig. S6 for mechanical characterisation).

### Mechanical Testing of Sensors within Implant Geometry Validates Simulation Results

Finite element models were produced to: 1) improve the sensor design by modelling electrode behaviour and the effect of reservoir dimensions and chip material on the sensitivity, and 2) obtain a theoretical force distribution in the trial insert upon application of an external force over a range of magnitudes and orientations relative to the cup, similar to a surgeon inserting and testing the implant during THR. The corresponding methods and results of the simulations are presented in the Supporting Information; here we focus on the experimental validation of the sensors.

To validate the simulation results, we used a custom-designed mechanical testing rig (Fig. 2a) to experimentally apply an external force from the head to the cup component in the hip implant. The rig consists of a polycarbonate (PC) support that houses a rotating brass stage. The stage has been machined to house the outer cup (Fig. 1g–i), which is held in place by semi-circular pegs. The sensors are inserted into the grooves between the inner and outer cups. The femoral head is attached to the moving component of a mechanical testing machine and is lowered into the cup to apply the desired forces. This rig was used with two mechanical testing machines: the Tinius Olsen 250 N and the Mayes 100 kN for lower and higher forces respectively. The stage can rotate and be locked in position to load the cup at angles of up to *θ* = 30° to the cup normal in approximately 5° increments.

**Figure 2.**
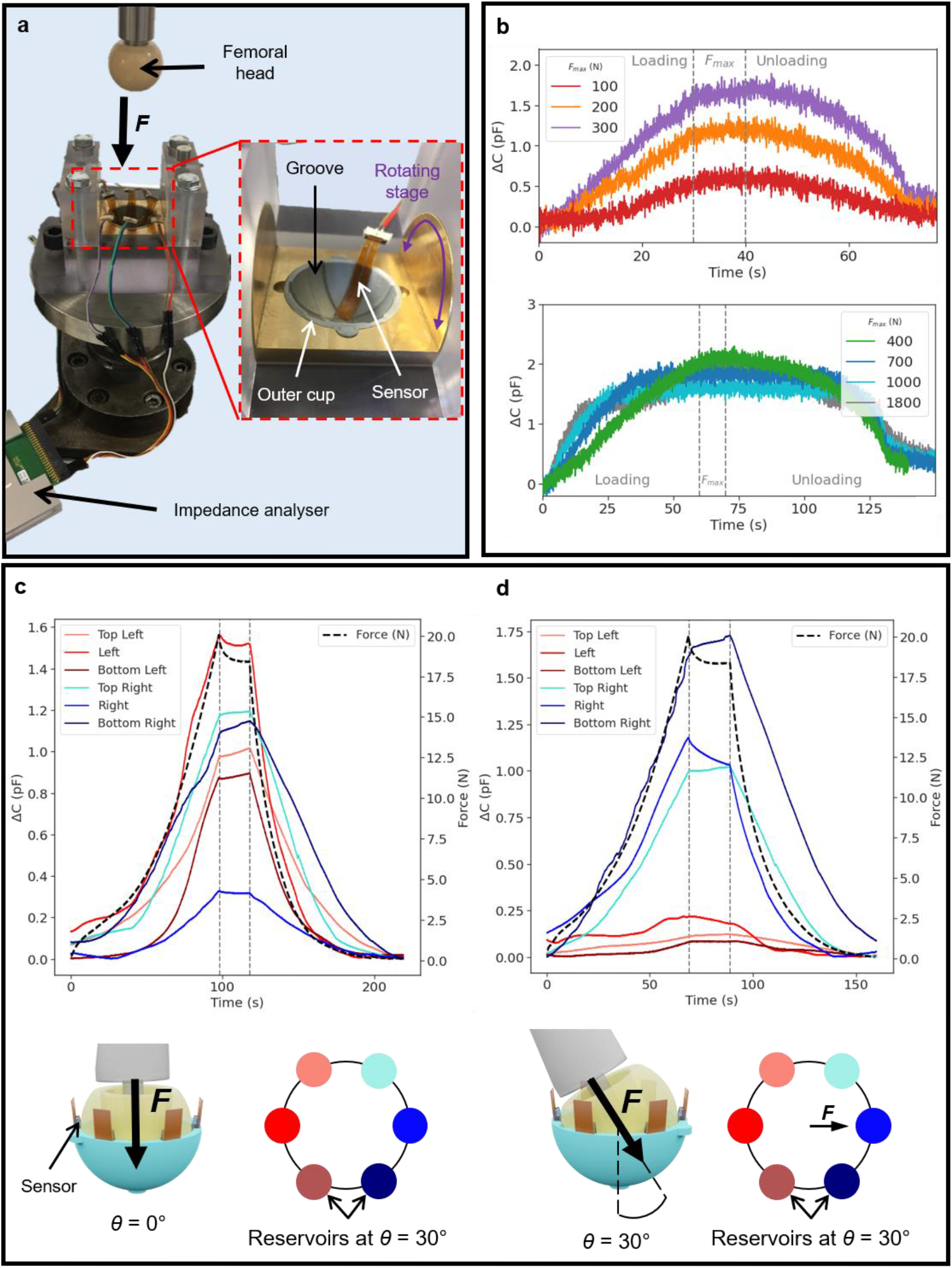
Mechanical testing of sensors in trial hip implant geometry. **a** Custom-built testing rig for use with the Mayes 100 kN Universal servohydraulic test machine. [Inset] The central stage rotates to simulate loading the insert cups at angles up to 30°. **b** Loading one sensor in the cup geometry up to a maximum applied force *F*_*max*_. [Top] There is a clear force-capacitance relationship between *F*_*max*_ = 100 N and 300 N. [Bottom] In order to reach higher forces, a different machine was used – therefore, the measurement duration is longer. The sensor is operational up to at least 400 N, above which the response begins to plateau. **c** Loading six sensors in the trial insert up to *F*_*max*_ = 20 N at a cup angle *θ* = 0°. The coloured lines and dots indicate different sensors. **d** Loading six sensors in the trial insert up to *F*_*max*_ = 20 N. The force is applied normal to a sensor’s reservoir, at a cup angle *θ* = 30°.

Fig. 2b demonstrates the effect of loading and unloading one sensor from applied force *F* = 0 N to a maximum force *F*_*max*_, which varied from 100 to 1800 N, over 14 loading cycles. During each cycle, the force was applied linearly from 0 N to *F*_*max*_, held at *F*_*max*_ for 10 seconds, then unloaded with the same linear relationship. A clear increase in capacitance values can be seen as *F*_*max*_ increases up to 400 N, above which the capacitance response starts to plateau, which would complicate calibration. The maximum capacitance change Δ*C* also decreases with increase in *F*_*max*_ above 700 N, indicating that fluid may be lost from the sensor. This is an appropriate force range for applications in hip surgery, but it can be increased by modifying the sensor design, for example by changing the channel and reservoir dimensions.

Fig. 2c shows the loading and unloading of six sensors incorporated into the cup, with *F*_*max*_ = 20 N and *θ* = 0°. The sensor reservoirs, the active sensing components, were located at 30° to the cup centre. The cup was loaded up to 20 N, the force was held at approximately 20 N for 20 seconds, then the cup was unloaded to 0 N. The sensors vary in response, which could indicate imbalance in the loading setup. Fig. 2d repeats this experiment for a cup angle of *θ* = 30°, showing the effect of the new cup orientation. The sensors with reservoirs closer to the axis of applied forces demonstrated a greater increase in capacitance. These experiments were repeated over 20 loading cycles without significant change in device performance, but the individual sensors can last up to at least 100 loading cycles without failure (see Supporting Information Fig. S15). The data in Fig. 2c and 2d also show a delay of a few seconds between applying the force and the sensors’ response, which may be due to force being applied too quickly relative to the fluid’s relaxation time.

Fig. 3 shows the calibration when a series of load cycles were applied, with the maximum force reached increasing from 50 N up to 200 N and decreasing back to 50 N. Fig. 3a demonstrates that the sensors can be loaded up to 200 N and then back to 50 N without a significant change in performance. Fig. 3b is a plot of the change in capacitance with force, calculated from capacitance-time and force-time data. The loading curves all show a similar sensitivity of 0.03 pF N^−1^, but the unloading curves show large hysteresis. This hysteresis is only present when calibrating the sensors inside the curved trial insert geometry – it is not present when the sensors are calibrated using the linear motor, when the sensors are flat^25^ (see Supporting Information Fig. S2). The hysteresis is consistent between cycles, so does not significantly affect the calibration. It may arise from the fluid wetting the channel’s internal surface, causing a time delay as the liquid returns to the reservoir. This hysteresis could be reduced by choosing a liquid that has less favourable interactions with the channel. Additional calibration, including varying the cup and stage angle and determining the consistency of samples, can be found in the Supporting Information Figs. S16 and S17.

**Figure 3.**
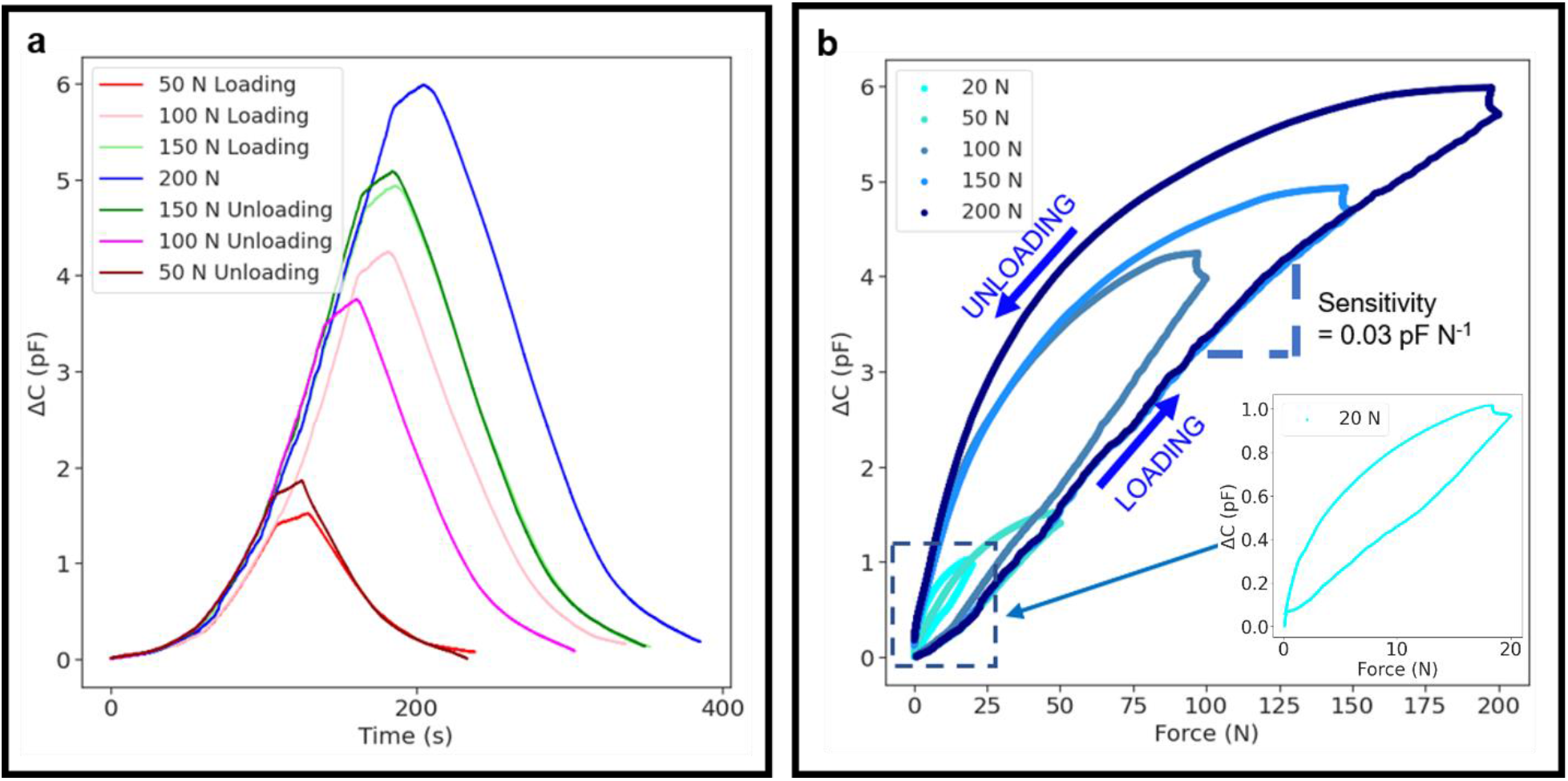
Sensor calibration and hysteresis behaviour in trial insert. **a** A series of loading curves showing change in capacitance over time, increasing *F*_*max*_ from 50 N to 200 N then decreasing it to 50 N. The experiments were performed in the order shown in the legend (top to bottom). **b** Force-capacitance calibration curves for the loading curves in **a**. The sensors exhibit significant hysteresis, with non-linear unloading curves indicating wetting of the channels by the fluid. This hysteresis is consistent between loading cycles, and therefore does not significantly affect calibration.

In this loading setup, the gradient of the capacitance-force calibration curve was lower – approximately 0.03 pF N^−1^ compared to 0.06 pF N^−1^ when calibrated outside of the cup^25^. It is therefore likely that the cup is acting to shield the sensors from these forces as previously indicated by the modelling results. This agrees with the observation that the maximum operating applied force for the sensors was considerably higher in this trial insert than when the sensors were calibrated on their own.

We have also characterised how modifications to both the chip design and material may affect device performance; such data (experimental and computational) is in the Supporting Information.

## Discussion

This paper summarises the design, production and characterisation of a next-generation hip implant technology that contains microfluidic force sensors, designed to replace part of the trial insert during surgery while the surgeon dynamically assesses the THR positioning, soft tissue tensioning, size and prosthetic impingement. This will give the surgeon quantitative data on the interfacial forces between the implant’s femoral head and insert during the THR, so that imbalance and improper positioning can be detected and corrected, decreasing the rate of implant failure. This is crucial, as surgeon experience is linked to fewer complications during surgery and better patient satisfaction post-surgery^28,29^.

Furthermore, it is well-documented that specialised hip surgeons are less likely to have complications as compared with general orthopaedic surgeons, and patients operated on by orthopaedic trainees report less satisfaction than surgeons with greater than 15 years of experience according to data from the Swedish Joint Registry^28–30^. The objective force feedback our technology provides has the potential to augment training and reduce learning curves in hip replacement surgery by allowing trainee surgeons to receive visual and haptic feedback of the accuracy of implant positioning during the trialling process.

Therefore, quantitative force data could be a useful tool to both improve the technique of experienced surgeons and as a training tool for non-specialists to perform the same surgery with the same outcomes.

Although accurate implant placement has now been addressed to an extent with navigation or robotic assisted techniques, difficulties with inappropriate soft tissue balance and prosthetic impingement remain unaddressed, leading to poor functional outcomes and accelerated wear^21^. Furthermore, they require a significant amount of investment and are not available for routine use in most hospitals worldwide^22^.

There are currently no commercial force sensors that are thin, conformable, and capable of bearing the large loads of a THR in a durable manner. The closest analogue is the VERASENSE (Orthosensor Inc., USA) for total knee replacement, which uses piezoelectric elements to quantitatively assess soft tissue balance and implant positioning to guide surgeons. However, this cannot be adapted for the hip as the hip joint contact zone is curved and has much less space than the knee.

Many different existing sensing mechanisms could potentially meet the clinical need for quantitative force sensing, including capacitive, resistive, piezoresistive, optical, triboelectric, magnetic, passive resonator and FET-based sensors^31^. Capacitive sensors have limited spatial resolution compared to resistive ones, but have a higher reliability^32–37^. Flexible microfluidic force sensors have been made previously for tactile and haptic sensing^38–40^, but could not reach the high forces required in this application. Some devices can reach higher resolutions, but each sensing element requires a large support system featuring electrical connections^41^ or power supplies.

Here, we have developed a capacitive microfluidic force sensor that is conformable and functional at the forces used in hip surgery. The sensors were experimentally and computationally validated for application in THR. Mechanical characterisation has demonstrated that the sensors can measure up to at least 400 N, which meets the expected requirements for the force range applied by the surgeon during THR^31,42^. This is orders of magnitude above the capability of similar microfluidic or flexible pressure sensors in the literature^43–45^. In the current design, the response beyond 400 N is saturated, but this can be easily adjusted by changing the chip and electrode geometry, the material stiffness and the trial insert design. The mechanical testing also revealed that there is no loss of sensor functionality when curved, and they can be repeatedly loaded at these high forces. The device currently incorporates six sensors, but improving the chip and electrode design, including wireless sensing, more sensing elements can be incorporated.

The sensors and trial implant provide minimal modification to the insert itself during surgery. Existing devices in the literature either modify the implant geometry^24,36,46^, which has been deemed expensive and technically challenging^31^, disturb the contact zone between the femoral head and acetabular cup^45,47^, or are implanted away from the contact area and therefore do not directly measure contact forces^48^.

Furthermore, the sensors and cups can be rapidly produced by a combination of aerosol jet printing and stereolithography 3D printing over a few hours, which is a significant improvement over traditional soft lithography methods that can be several times more expensive and time-consuming^49^. Therefore, while these sensors have been demonstrated to work in the ball-and-cup hip joint geometry, there is scope to adapt the sensor design to work in other areas of the body, including the knee, in the patellofemoral joint and in medial and lateral compartmental replacement, as well as in the spine, shoulder and ankle.

While these sensors have advantages of flexibility, greater force range and ease of manufacture, they must also be sensitive enough to detect imbalance. Studies using sensor-guided technology in the knee^50,51^ that used two force-sensing components defined that the implant is ‘balanced’ if the two sensing components registered a difference of 67 N, which our sensors can easily distinguish. In the future, *in vivo* testing will be carried out to demonstrate the technology’s usefulness in desired settings. Biocompatibility tests will be performed to determine the host’s response to the sensors. Cadaveric testing will trial the usage of the trial insert and are to be performed imminently. The sensors have been tested and shown to work at ambient temperatures (10-50 °C).

In summary, we present a thin, flexible, and robust capacitive microfluidic force sensor, that can be fabricated quickly and cheaply using additive manufacturing. A custom-made mechanical testing rig was designed, produced, and used to demonstrate that the sensors can be loaded up to the forces typically applied by the surgeon during THR. Due to the sensor design’s customisability and adaptability, in the future these sensors could be used in a range of different applications where force monitoring and balancing is essential.

## Methods

### Sensor Fabrication

#### Flexible Resin microfluidic chip fabrication

The CAD file for the chip was custom-designed using Creo Parametric 6.0 (PTC, USA), and printed using on the Formlabs Form 3B stereolithography (SLA) 3D printer (Formlabs, USA) using the resin’s smallest layer thickness of 50 μm. After printing, the chip was then washed in isopropanol (IPA) for 20 minutes in a sonicating bath before being transferred to the oven (FormCure, Formlabs, USA) for curing at 60 °C for 10 minutes. The supports were then removed.

#### Interdigitated electrodes fabrication

1-1.5 mL silver ink was produced by diluting Novacentrix Ag nanoparticles aerosol ink (JS-A221AE) 1:3 by volume with DI water. The ultrasonic atomiser in the aerosol jet printer (AJP) was used with a tip size of 150 μm to print the electrodes on a 75 μm-thick Kapton (PI) film (RS Components Ltd., UK). Ink and sheath flow rates were 20 and 80 sccm respectively. The overall dimensions of the electrodes are 20 mm x 0.5 mm. Silver connecting pads were printed at the end of the electrodes, which fit a 2-pin flexible printed circuit (FPC) connector. The silver was then cured in the oven at 150 °C for 2 hours. To print the insulating polyimide (PI) layer on top of the electrodes, a 15 mL PI ink made from a mix of poly(pyromellitic dianhydrideco-4,4’-oxydianiline), amic acid solution (12.8 wt.% in 80% NMP / 20% aromatic hydrocarbon, Sigma-Aldrich) and N-methyl-2-pyrrolidone (NMP, Sigma-Aldrich) at a 1:1 volume ratio was used with the pneumatic atomiser and a tip size of 300 μm. All the electrodes were covered with PI except for the connecting pads. To cure the polyimide, the electrodes were placed in the oven at 200 °C for 2 hours. The Kapton was then cut into the correct shape and size.

#### Bonding the microfluidic chip to the electrode layer^25^

The adhesive layer connecting the microfluidic chip and Kapton electrode layer was made from laser-cut double-sided tape (RS Components Ltd., UK). The layer was designed in AutoCAD 2020, and then cut into the tape using a laser cutter (Epilog Zing 16, 30 W). Vector mode was used, with a resolution of 500 DPI and a frequency of 2500 Hz. The vector speed and power were 90% and 10% respectively. One side of the tape was then adhered to the microfluidic chip. The chip was then pressed onto the Kapton layer with the adhesive layer in between. The alignment between the chip and Kapton was then checked under an optical microscope. To minimise air bubbles between the chip and Kapton layers, a weight was put on top of the chip layer to press the two layers as the glue cured. The glue was then left to cure for a few hours.

#### Fluid injection

Once the glue has cured, fluid was injected into the channel to completely fill the reservoir up to the entrance to the channel. The injection hole is located adjacent to the reservoir but at the opposite end to the channel and was incorporated into the CAD file of the chip. The fluid, a 2:1 by volume glycerol-water mixture, was injected until it just filled the reservoir. The ratio was chosen to obtain a balance between a low volatility and high dielectric constant. The hole was then sealed with the silicone sealant to prevent the fluid evaporating.

### Experimental Measurement of Sensor Response

#### Impedance Analyser

The Sciospec ISX-3 was used to measure the electrical impedance of the sensors. By using the simple circuit model for the sensor as a resistor in series with a capacitor, the impedance *Z* can be converted into a capacitance *C* using the equation *C* = −(2*πf* . *Im* {*Z*})^−1^ where *f* is the measurement frequency in Hz. To determine *f*, frequency sweeps were conducted from 1 kHz to 1 MHz. Frequencies below 20 kHz were determined to have high noise, whereas higher frequencies had low impedance magnitudes. A compromise frequency of 500 kHz was selected for further measurements for a good signal-to-noise ratio. Three measurements were carried out at each applied force and averaged over three different loading cycles.

#### Linear Motor and Load Cell

A linear motor (LinMot, Switzerland) with a 20 N load cell (PCM Ltd., UK) was also used to test the sensors individually before incorporation into the implant geometry. It was found in practice that the load cell could read forces up to 40 N if a sufficient cooling system was put in place to prevent overheating.

#### Mechanical testing

Mayes 100 kN and Tinius Olsen 250 N mechanical machines were used to test the *in vitro* response of the sensors in the custom-made testing rig. To determine the maximum forces at which the sensor calibration broke down (for example, when the capacitance response plateaued), the femoral component was attached onto the load cell of the Mayes 100 kN servohydraulic testing machine, while the custom testing rig containing the acetabular component was screwed into the machine itself. For calibration, the testing rig was attached to the Tinius Olsen mechanical testing machine with a 250 N load cell. Details on the mechanical testing of the bulk microfluidic chip material are included in the Supporting Information.

## Acknowledgements

L.I. acknowledges support from an EPSRC Doctoral Training Partnership studentship (EP/R513180/1). T.W. acknowledges support from an EPSRC Doctoral Training Partnership studentship (EP/T517847/1). S.K.-N. is grateful for support from ERC Starting Grant (Grant No. ERC-2014-STG-639526, NANOGEN). S.K.-N. and Q.J. acknowledge support from the Centre of Advanced Materials for Integrated Energy Systems ‘‘CAM-IES’’ grant EP/P007767/1. J.C. is currently supported by a Wellcome Trust Institutional Strategic Support Award to the University of Exeter (204909/Z/16/Z); for the purpose of Open Access, the author has applied a CC BY public copyright license to any Author Accepted Manuscript version arising from this submission.

## Competing interests

L.I., Q.J., V.K., J.C., and S.K.-N. are named inventors on a patent application by Cambridge Enterprise that covers the technology described in this study. S.K-N, V.K. and J.C. are co-founders and directors of ArtioSense Ltd., incorporated to translate and commercialise aspects of this work. VK is Consultant for Arthrex and Smith and Nephew Limited.

