## Supporting Information for "Conformable and robust force sensors to enable precision joint replacement surgery"

### Methods

#### *Experimental Mechanical Characterisation of Sensor and Trial Insert Materials*

Examples of candidate materials for the microfluidic chip include polydimethylsiloxane (PDMS) and "Flexible Resin" (Formlabs, USA), a stereolithography (SLA) 3D printing resin. Flexible Resin has a larger hardness (Shore A hardness = 80-85[1]) than PDMS (approx. Shore A hardness = 44 at room temperature[2]) and is therefore likely to be stiffer. Formlabs do not reveal the exact composition of Flexible Resin, so mechanical characterisation was done to determine its suitability over PDMS.

While PDMS is one of the materials used to construct the channel and reservoir in this paper, it has many limitations for this application. It absorbs small molecules, decreasing the volume of fluid in the reservoir and making the sensor more susceptible to air bubble intrusion[3–5]. There is some variability between the mechanical properties of different batches of PDMS, as it heavily depends on the extent of mixing that occurs, the curing temperature and curing duration. Therefore, some attempt has been made to find suitable alternatives to PDMS. The desired criteria are for a material that (i) can withstand forces up to similar loads applied by surgeons, estimated to be in the range of hundreds of Newtons, (ii) has an appropriate stiffness such that at these forces the reservoir is mostly depleted (for maximum sensitivity) and (iii) has a scalable manufacturing process.

Some candidate materials are available in the library of Engineering Resins available from Formlabs for their SLA printers (Formlabs, USA). SLA printing is already established to have good scalability and high resolution[6]. Some possible resins include the Flexible, Elastic, Durable and Tough resins. The exact chemical compositions are unknown due to their proprietary nature, but some mechanical properties have been provided by Formlabs, who have suggested materials that are similar to these resins (PMMA is similar to Grey Resin, for example).

The mechanical properties of PDMS, Flexible Resin and Durable Resin were characterised. Dog-bone shaped samples were used for tensile testing, with dimensions corresponding to the American Society for Testing and Materials (ASTM) standard D412-C, and were prepared in a laser-cut poly(methyl methacrylate) mold (for PDMS) or direct stereolithography 3D printing using the Formlabs Form 3 3D printer (for Flexible Resin, Durable Resin). Compressive testing samples consisted of 10 mm diameter, 10 mm height cylinders, as in ASTM D575-91[7], and were molded in a drilled PMMA sheet (for PDMS) or SLA printed (for Flexible Resin and Durable Resin). For the PDMS samples, PDMS (Sylgard 184, Dow, USA) was mixed with curing agent in a 10:1 ratio by weight, poured into the molds and left to cure in an oven (Heratherm OGH60, Thermo Fisher, USA) at 70 °C for 2 hours. Once cured, the PDMS was removed from the mold.

Force-extension curves were measured with an extension-controlled universal tester (Tinius Olsen 5ST, UK) equipped with a 250 N load cell. For the PDMS samples, the extension speeds were 10 mm $min^{-1}$ in tension and 5 mm $min^{-1}$ for the smaller compression samples. For Flexible Resin, the extension speeds were 20 mm $min^{-1}$ in tension and 3 mm $min^{-1}$ in compression. For Durable Resin, the extension speeds were 20 mm $min^{-1}$ in tension and 3 mm $min^{-1}$ in compression. These rates, lower than suggested by testing standards[7,8], were chosen to be more representative of the strain rate applied on the sensors described in this paper. Eccentric roller grips and 5 cm diameter compression plates were used to fix samples.

Non-linear least-squares fitting was then carried out in COMSOL’s Optimisation module and Microsoft Excel. A cost function, evaluating the difference between measured and fitted curves, was minimised to identify optimal parameters for the Neo-Hookean, Mooney-Rivlin, Ogden and Yeoh hyperelastic models[9].

### *Finite Element Modelling of PDMS-Based Sensors*

#### Optimisation of Sensor Design

To increase the range of forces that the sensors can measure, a variety of modifications were made to the design of the sensor, including the reservoir width and height, the channel height, the number of columns and the PDMS curing ratio. These were compared with experimental calibration data.

Finite Element Modelling (FEM) was conducted using COMSOL Multiphysics 5.4 (COMSOL Inc., USA), a Finite Element Analysis (FEA) software, to optimise the sensor design before manufacture, to improve both the device sensitivity and increase the range of forces to achieve the 450 N target. The sensor was not modelled as a single model, as this would require analysis of fluid dynamics and electrostatics in response to mechanical deformation, which is both complex to model and computationally demanding. Instead, three models were produced to simulate the key components of the sensor operation:

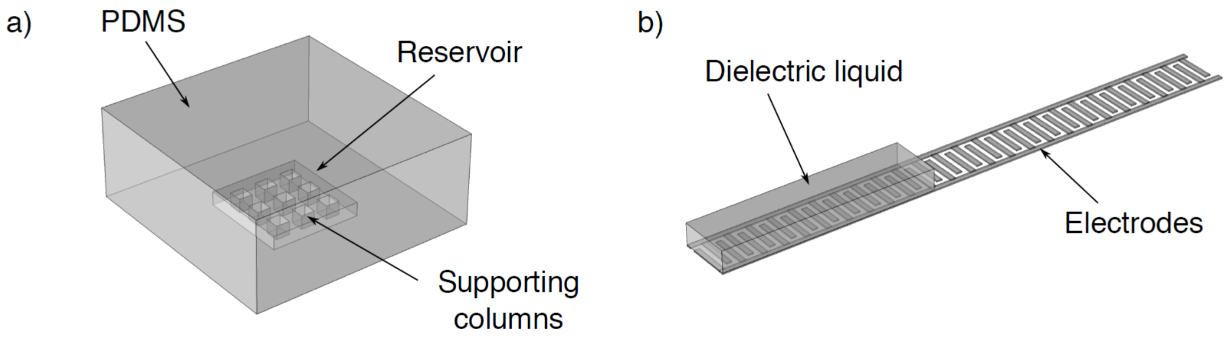


**Figure S1 | Visualisation of studied FEM geometries.** (a) The reservoir model, which consists of a fluid reservoir (Length x Width x Height: 2 mm x 2 mm x 0.3 mm) embedded in a soft elastomer such as PDMS (5 mm x 5 mm exposed area). (b) The electrode model (20 mm electrode length, 0.1 mm electrode width and separation, 0.5 mm x 0.2 mm channel cross-section).

Two models were produced to simulate the response of an individual sensor to an applied load. The *electrode model* simulates the response of the electrodes (including capacitance values) when several parameters are varied, including the length of the liquid covering the electrodes, the shape of the electrodes and the type of liquid. The length of fluid displaced along the channel is converted into a capacitance value. The *reservoir deformation model* simulates the deformation of the reservoir as a force is applied. We model the effects of different reservoir geometries, varying the width, height, number and shape of columns, reservoir material, and PDMS curing ratio. The applied force is converted into a change in internal volume of the reservoir, which in turn is equivalent to the volume of fluid displaced along the channel. To simulate the deformation of PDMS, the Mooney-Rivlin 5-parameter hyperelastic model was considered most suitable for the range of strains obtained in the model[26] (see Figs. S5 and S5 for the fitting of hyperelastic deformation models to the microfluidic chip materials). Modelling the sensor response in planar geometries was covered in the group's previous work[25] – in this study, the sensor modelling is conducted using the curved implant geometry.

**Reservoir model (Fig. S1a):** To model the deformation of the reservoir as a force is applied, in particular calculating the change in volume of the reservoir (and therefore, of the fluid). The effects of different reservoir geometries were calculated, including varying the width, height, number and shape of columns, and reservoir material (including PDMS curing ratio). The effect of curvature was also considered, resembling the insertion of the sensor in the implant geometry. Simulations were performed using COMSOL's Structural Mechanics module. For this model, the volume was calculated by integrating the internal surfaces of the reservoir. The boundary conditions were a compressive force on the top surface and a fixed bottom surface, although the sides were also fixed in one simulation to determine the effect of enclosing the sensors in the grooves. The PDMS itself was modelled as a hyperelastic material, using the Mooney-Rivlin 5 parameter model, with the parameter obtained from mechanical testing (see Table S1). A free tetrahedral mesh was applied - the element sizes were generally around 0.25 mm, but the reservoir boundary concentrated more stress and so was meshed more finely at 0.09 mm.

**Electrode model (Fig. S1b):** The reservoir model outputs a change in volume of the reservoir under applied stress, which is equivalent to a volume of displaced dielectric liquid. The electrode model then determines the change in capacitance when this volume of fluid covers the interdigitated electrodes, resulting in a relationship between the applied force and the capacitance of the sensor. This was conducted using COMSOL's Electrostatics module. In this case, the boundary conditions were grounding one of the electrodes while raising the other to a potential of 1 V, and grounding the surrounding environment. The free tetrahedral mesh had a small element size of 0.0075 mm for the electrodes. The effect of curving the electrodes to the radius of the acetabular cup was determined.

#### Simulation of Sensors in Trial Insert

A third finite element model was produced, to simulate the force distribution in the trial insert and therefore predict theoretical relative force readings between sensors located in different areas of the cup.

**Ball and Socket model (Fig. S14):** This was designed to model the ball and cup deformation, including the applied forces at different points in the cup when a certain force is applied by the head. The femoral head was made from a truncated sphere of alumina of radius 15 mm, using COMSOL's parameters for alumina (Young's Modulus $E$ = 300 GPa, Poisson ratio $\nu$ = 0.22). The acetabular cup was modelled as a 1.5 mm-thick hemisphere with the same internal radius as the femoral head, made from HDPE because of its similarity to Formlabs' Durable Resin. The orientation of the head with respect to the cup axis ($z$-axis) was described by two angles $\theta_1$ and $\theta_2$ about orthogonal rotation axes ($x$- and $y$-axes, respectively). COMSOL's Structural Mechanics module was used to implement this. A *contact pair* was established between the head and cup's outer surface. The femoral head was set as the source and the cup was set as the destination, following recommendations[52]. The model was solved using the *penalty* method. The boundary conditions were that the external cup surface was fixed, and that the femoral head was displaced down its axis (at angles of $\theta_1$, $\theta_2$ to the cup's $z$ axis as described earlier). Friction between the head and cup was neglected. The femoral head was meshed coarsely, from 2 mm elements. Both elements of the contact pair, the internal cup surface and femoral head surface were meshed as 0.6 mm and 1.1 mm respectively.

#### *PDMS-Based Sensor Fabrication*

The PDMS-based sensors were manufactured in a different way to those made from Flexible Resin, in that the microfluidic chip was made by pouring PDMS into a 3D-printed mould:

**PDMS microfluidic chip fabrication:** a mold was printed using an SLA 3D printer (Form 3B, Formlabs, USA) using Formlabs' proprietary Grey resin. The CAD file for the molds was custom-designed using Creo Parametric (PTC, USA), and the smallest layer thickness of 25 μm was chosen. After printing, the mold was washed in isopropanol in a sonicating bath for 15 minutes to remove excess resin. The mold was then cured in an oven (FormCure, Formlabs, USA) at 60 °C for one hour. PDMS (Sylgard 184, Dow, USA) was mixed with curing agent in a 10:1 ratio by weight, poured into the mold and left to cure in an oven (Heratherm OGH60, Thermo Fisher, USA) at 70 °C for 2 hours. Once cured, the PDMS was removed from the mold and cut into the appropriate size.

**Bonding PDMS to Kapton[12]:** a thin layer of primer (DOWSIL PR-1200 RTV Prime Coat, Dow, USA) was first applied on the surface of the Kapton film and left for 1 hour until fully dried. A thin layer of silicone sealant (DOWSIL 3140, Dow, USA) was applied on top of the primer, taking care not to cover the electrodes, and then attached to the PDMS microfluidic chip. The alignment between the PDMS and Kapton was then checked under an optical microscope. In order to minimise air bubbles between the PDMS and Kapton layers, a weight was put on top of the PDMS layer to press the two layers as the glue cured. The glue was then left to cure for a few hours.

**Fluid injection:** Once the glue has cured, a hole was made in the PDMS layer using a syringe. The hole is located adjacent to the reservoir but at the opposite end to the channel. The fluid, a 2:1 by volume glycerol-water mixture, was injected until it just filled the reservoir. The ratio was chosen in order to obtain a balance between a low volatility and high dielectric constant. The hole was then resealed with the silicone sealant.

The fabrication of the electrode layer is unchanged from the Flexible Resin sensors, and thus its fabrication details are given in the main manuscript.

#### *Experimental Characterisation and Optimisation of PDMS-Based Sensor Design*

#### Calibration and Fatigue Properties

Capacitance-force calibration was conducted using a linear motor (LinMot, Switzerland), the setup of which can be seen in Fig. S3. The pressing arm was made from a flat-head screw with a nitrile rubber sheet wrapped around it. The sensor was then attached between the pressing arm and the load cell. To apply load to the sensor, the linear motor was programmed to advance the pressing arm in intervals of 100 μm. To calibrate the sensor, forces read from the load cell were recorded while the impedance of the electrodes was monitored by an impedance analyser. The load cell was rated to read forces up to 20N, although it was found in practice that this could go up to 40N if a sufficient cooling system was put in place to stop it from overheating.

In order to determine the optimal design for the microfluidic chip, a variety of chip parameters were varied, including: channel dimensions; reservoir dimensions and shape; the presence, size and cross-sectional shape of supporting columns; and PDMS curing ratio. Each of the sensors with a different design was calibrated using the linear motor up to 20 N.

In addition, the fatigue properties of a typical sensor were investigated. An oscillating force between 0 - 11.67 N was applied to a PDMS-based sensor using the linear motor, with the pressing arm speed set at 0.584 mm $s^{-1}$, with the aim to reach 100 loading cycles before significant change in device performance.

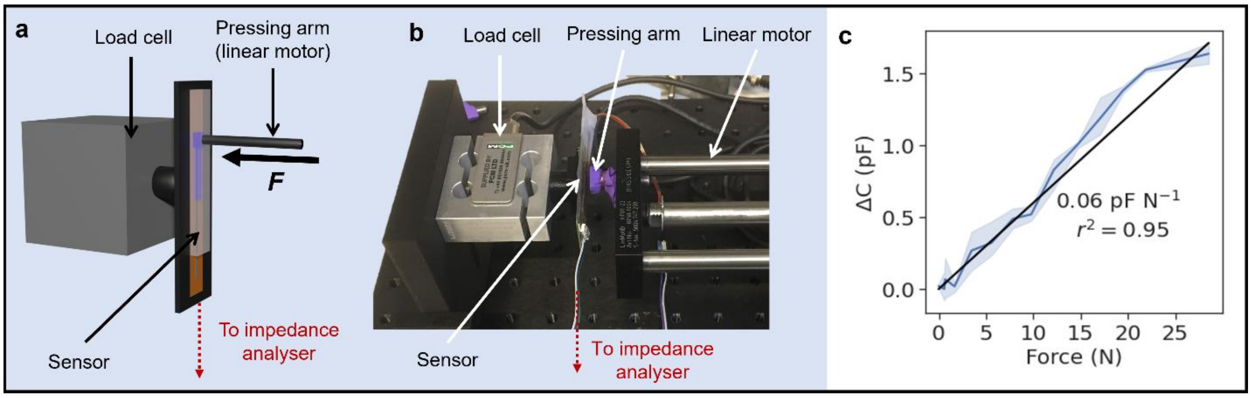


**Figure S2 | Calibration of sensors using a linear motor. a** Schematic diagram of linear motor operation – a pressing arm exerts a force on the sensor, which is adhered to a load cell to measure the applied force. The sensor impedance is measured with an impedance analyser. **b** Photograph showing one sensor in the linear motor setup. **c** Typical calibration of a Flexible Resin-based sensor using the linear motor.

### Results

#### *Experimental Mechanical Characterisation of Sensor and Insert Materials*

##### PDMS

**Effect of PDMS curing ratio (Fig. S3):** A lower curing ratio of PDMS led to slightly decreased sensitivity, both in simulations and experiments. Surprisingly, the mechanical properties of 5:1 and 10:1 PDMS were similar in compression, leading to similar responses of sensors made from 5:1 and 10:1 ratio PDMS.

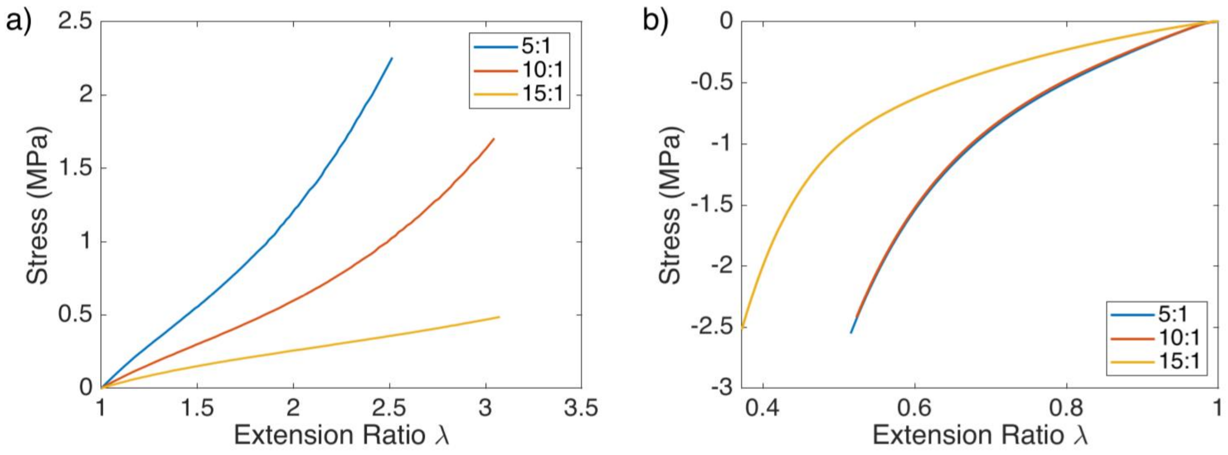


**Figure S3 | Experimentally measured mechanical properties of PDMS at different curing ratios. a** Uniaxial tension, carried out to failure. **b** Compression, terminated near the limit of the load cell range at 200 N.

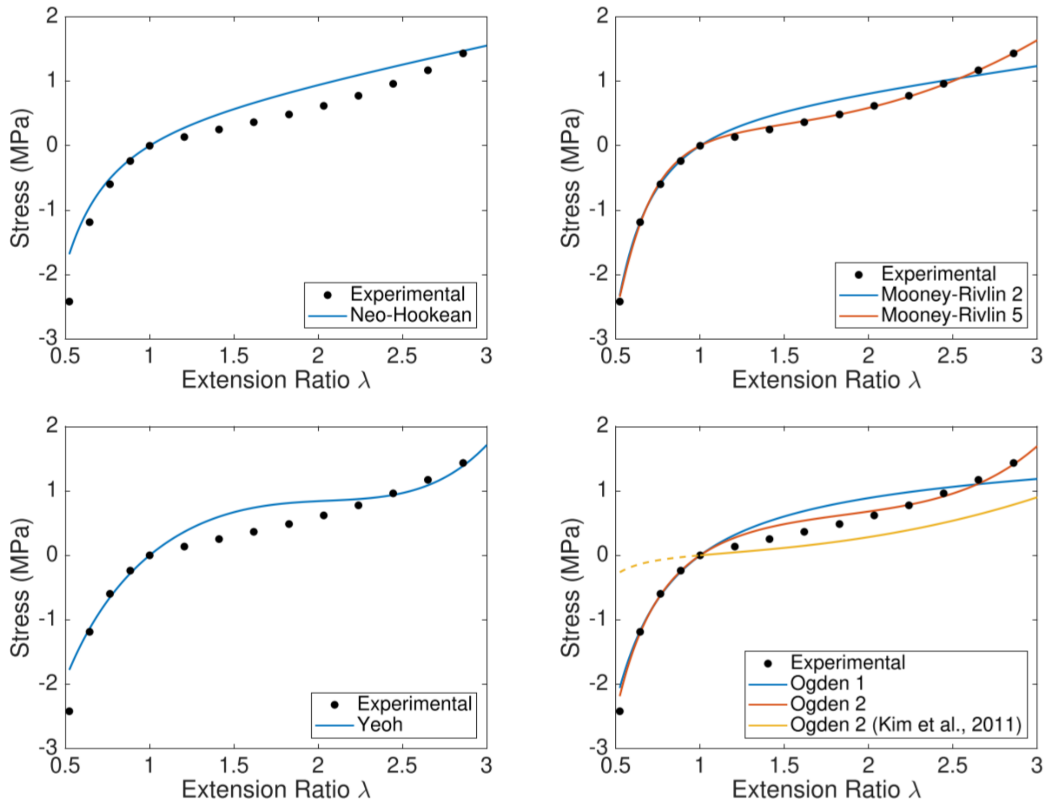

**Figure S4 | Hyperelastic models fitted to the experimental stress-extension ratio curve for 10:1 PDMS.** For comparison, the best-fit numerical curve from Kim *et al.* (2011)[10], Ogden 2nd order, is included and extrapolated in compression.

**Table S1 | Reduced chi-squared values for different hyperelastic models fitted to measured stress-extension ratio curves.** Values significantly greater than 1 indicate poor fit whereas values smaller than 1 show overfitting with respect to experimental error.

| **Reduced chi-squared, $\chi_R^2$** | **PDMS Curing Ratio** | | |
|---|---|---|---|
| | **5:1** | **10:1** | **15:1** |
| Neo-Hookean | 22.73 | 7.24 | 480.0 |
| Mooney-Rivlin, 2 parameters | 23.61 | 2.83 | 30.75 |
| Mooney-Rivlin, 5 parameters | 3.20 | 0.96 | 47.67 |
| Yeoh | 12.33 | 8.59 | 546.7 |
| Ogden, 1st order | 17.96 | 5.24 | 136.0 |
| Ogden, 2nd order | 5.67 | 2.67 | 96.40 |

### Flexible Resin

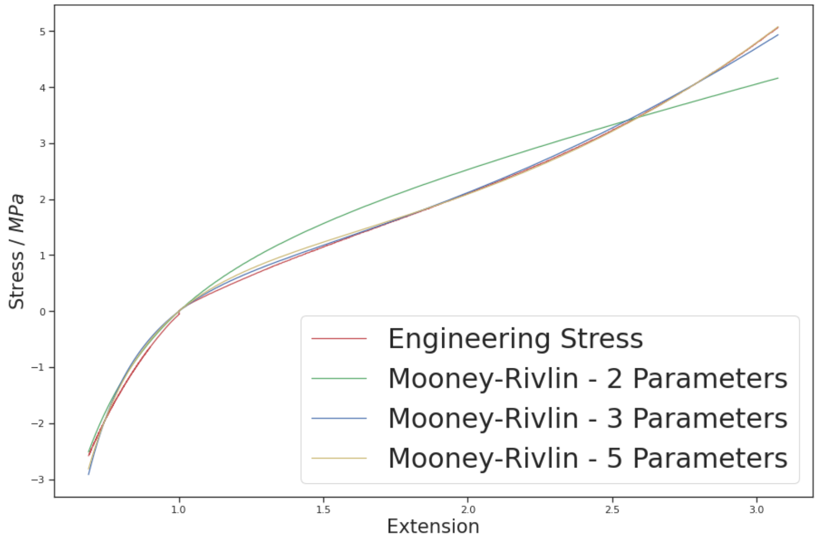


**Figure S5 | Mooney-Rivlin hyperelastic models fitted to the experimental stress-extension ratio curve for Flexible Resin.**

The calculated hyperelastic parameters from optimisation are given in Table S2:

**Table S2 | Calculated Mooney-Rivlin parameters that provide the best fit for the stress-extension data for Flexible Resin.**

| | **Mooney-Rivlin 2-Parameter** | **Mooney-Rivlin 3-Parameter** | **Mooney-Rivlin 5-Parameter** |
|---|---|---|---|
| **Parameter** | Value / MPa | Value / MPa | Value / MPa |
| $C_{10}$ | 0.62691 | 0.10901 | 0.19850 |
| $C_{01}$ | 0.36902 | 0.58551 | 0.58126 |
| $C_{11}$ | N/A | 0.09690 | 0.00001 |
| $C_{20}$ | N/A | N/A | 0.03392 |
| $C_{02}$ | N/A | N/A | 0.00000 |

#### Durable Resin

Using the linear region of the stress-strain curve in Fig. S4, the Young's modulus of Durable Resin is approximately 162 MPa, although other sources indicate a much higher modulus of 1.0 GPa[11].

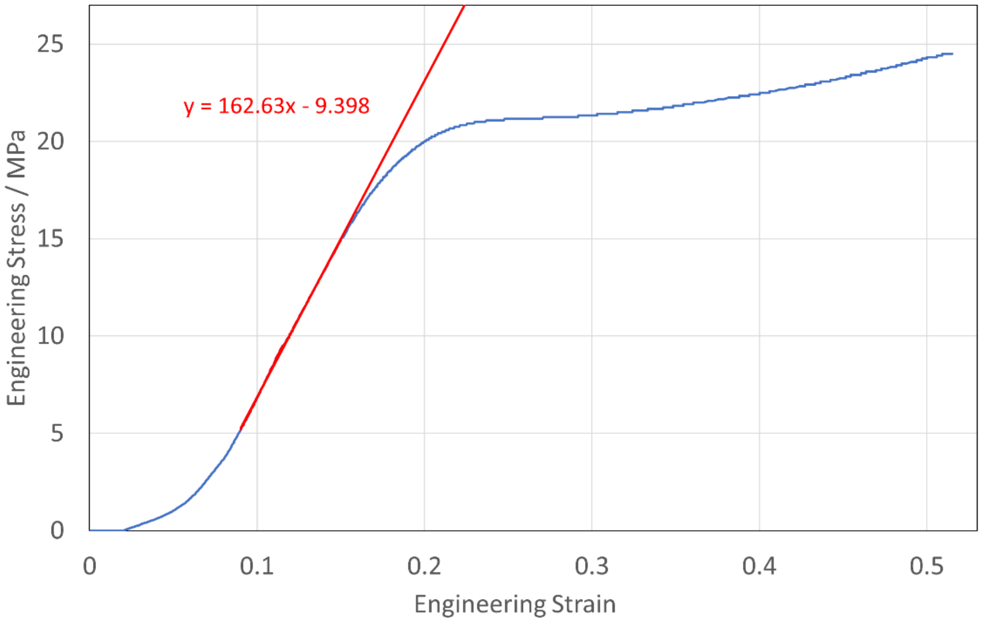


**Figure S6 | Engineering stress-strain curve for Durable Resin.** The behaviour is non-linear at very low and high strains, with an approximately linear region in-between.

### *Finite Element Modelling of PDMS-Based Sensors*

#### Optimisation of Sensor Design

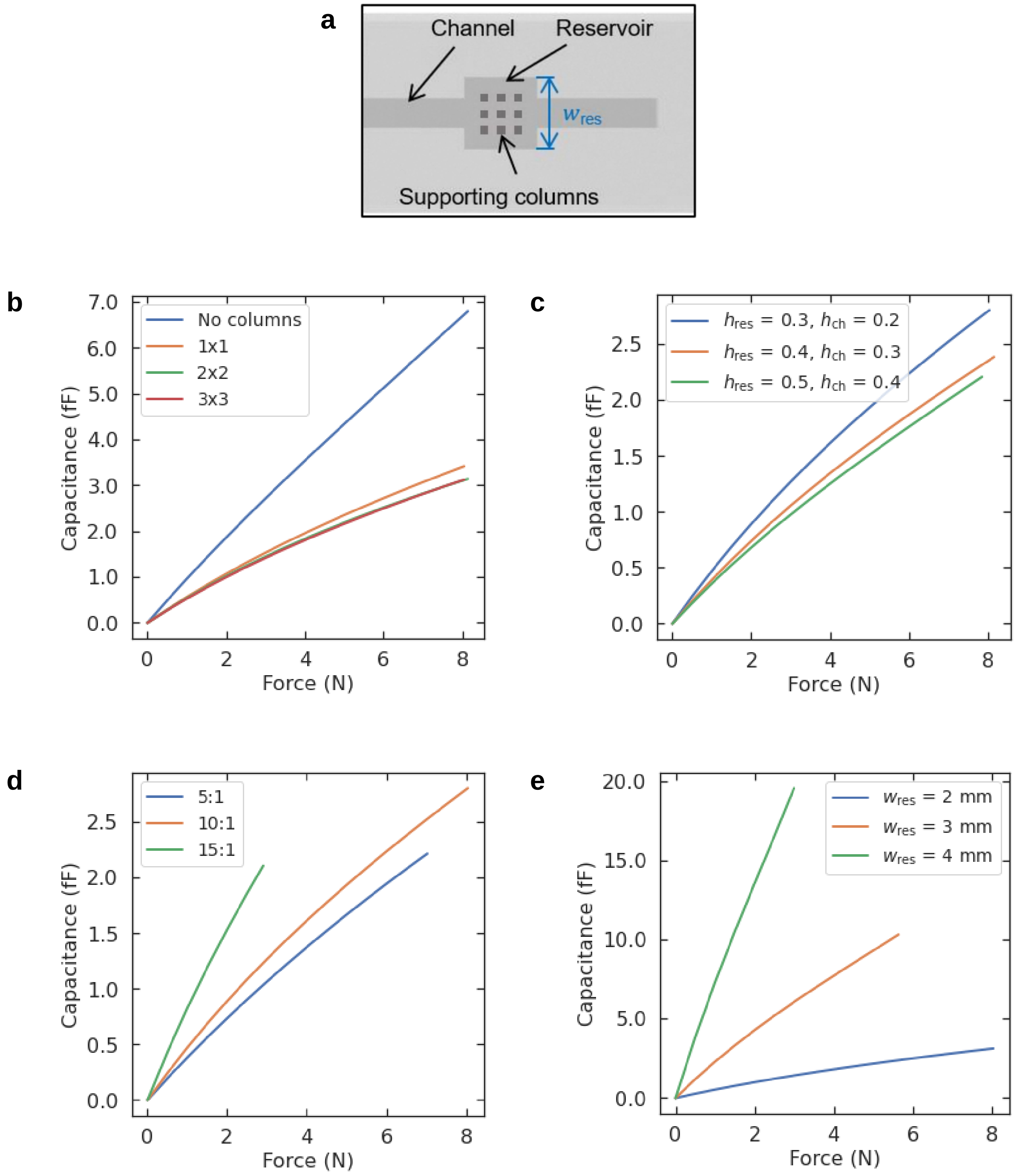


**Figure S7 | Effect of changing reservoir design on sensor sensitivity, for a reservoir supported by square cross-section columns.** For most simulations, the model could not converge past 8 N. For other simulations, convergence failed at lower forces. **a** Schematic of the reservoir, showing the supporting columns. **b** Effect of varying the number of columns. **c** Effect of varying the reservoir height $h_{res}$ and channel height $h_{ch}$. **d** Effect of varying the ratio of PDMS base to curing agent (by mass). **e** Effect of varying the width of the square reservoir, $w_{res}$.

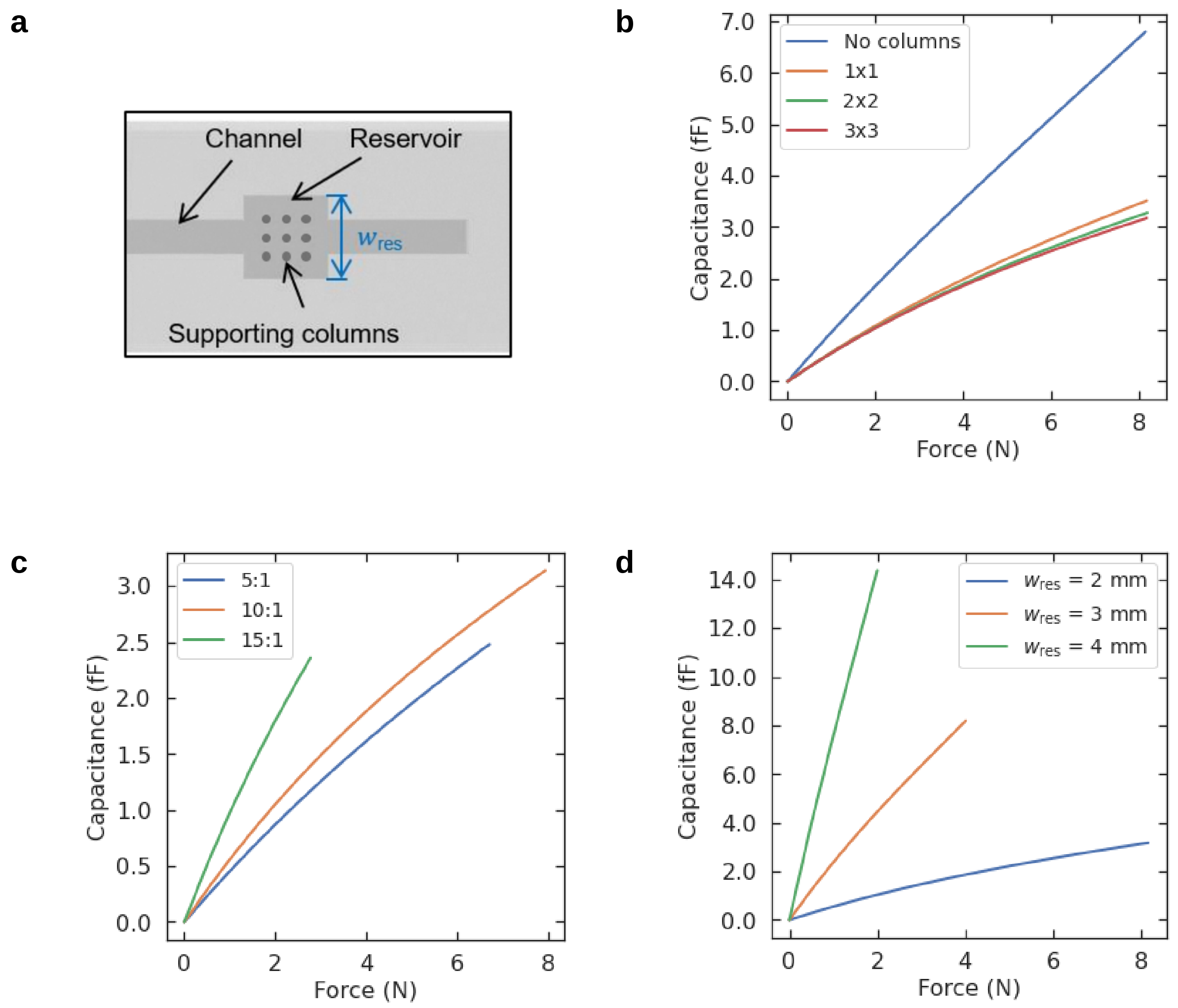


**Figure S8 | Effect of changing reservoir design on sensor sensitivity, for a reservoir supported by circular cross-section columns.** For most simulations, the model could not converge past 8 N. For other simulations, convergence failed at lower forces. **a** Schematic of the reservoir, showing the supporting columns. **b** Effect of varying the number of columns. **c** Effect of varying the PDMS curing ratio**.** **d** Effect of varying the width of the square reservoir, $w_{\mathrm{res}}$.

Figs. S7-S10 indicate that the cross-sectional shape of the supporting columns has very little effect on the sensitivity. The most important feature is the volume of liquid in the reservoir that can be displaced into the channel, which depends on properties such as the number of columns and reservoir width and height.

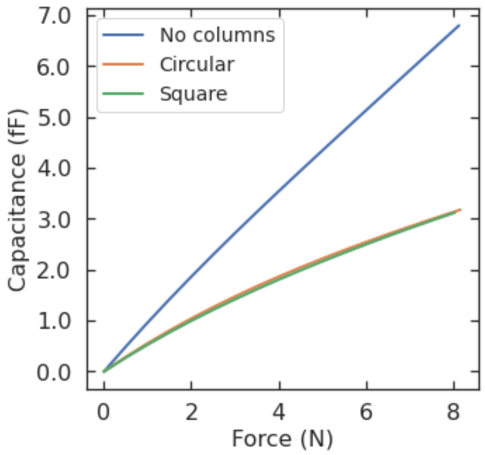


**Figure S9 | The effect of changing the cross-sectional shape of the supporting columns, showing the effect of having square columns, circular columns, or no columns at all.** Note that as per Figures S6 and S7, the column cross-sectional shape has little effect.

The effect of changing the shape of the supporting columns, or removing the columns entirely, is shown in Fig. S9. PDMS with a 10:1 curing ratio was assigned as the microfluidic chip material. The reservoir had a square cross-section ($w_{\mathrm{res}}$ = 2 mm) and the supporting columns had either square or circular cross sections, or were absent entirely. Removing the columns decreases the complexity of fabrication, as it is easier to fill the channel with fluid by avoiding air bubbles, but is also less computationally complex to simulate. The disadvantage to removing the columns for PDMS-based devices is the mechanical stability– the columns prevented the reservoir from collapsing in on itself when high forces were applied. Due to the increased stiffness, Flexible Resin-based devices did not have this issue and therefore the columns were not used.

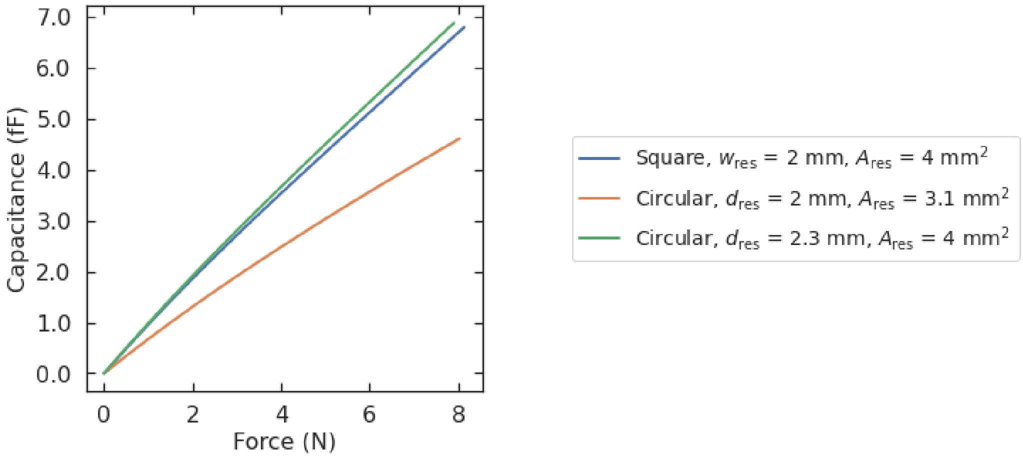


**Figure S10 | Showing the effect of changing the reservoir shape on the sensitivity.** This figure compares the effect of a square cross-section with width $w_{\mathrm{res}}$ = 2 mm (blue), a circular cross-section with a diameter $d_{\mathrm{res}}$ = 2 mm (orange), and a circular cross section with the same cross-sectional area $A_{\mathrm{res}}$ as the square reservoir (green). The two simulations with the same reservoir cross-sectional area have similar responses. The reservoirs in this simulation have no supporting columns.

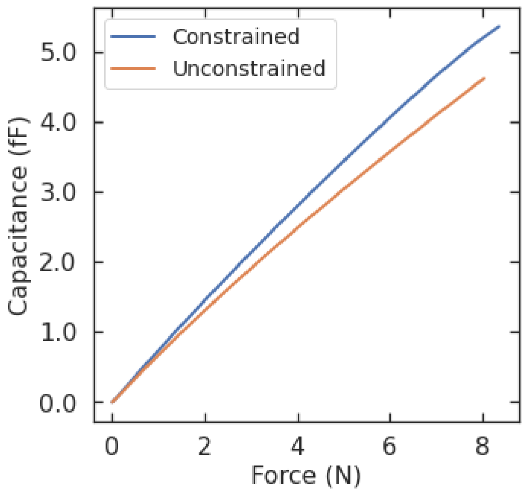


**Figure S11 | Applying a lateral constraint to the deforming reservoir**, to simulate the sensor being 'constrained' in the cup grooves.

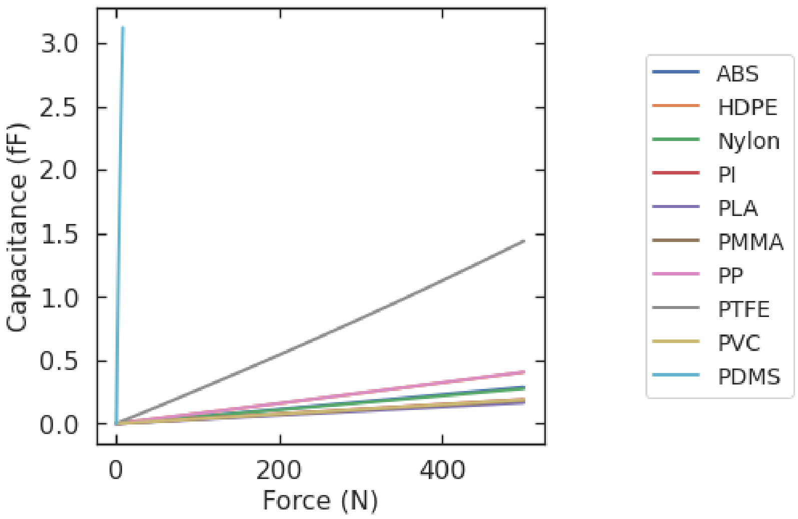


**Figure S12 | Brief comparison of PDMS to other common polymers.** PDMS is simulated as a hyperelastic material, the others are treated as linear elastic. Due to its relatively low stiffness, a PDMS sensor would exhibit a greater fluid displacement per unit force, leading to a larger capacitance change for low forces.

In addition, 15 mm curvature was introduced to both the reservoir and electrode models to determine the effect of curvature on sensor response. In the electrode model, the sensitivity increased by less than 0.5%, from 9.43 to 9.46 pF $\mu L^{-1}$ (negligible compared to other factors e.g. choice of mesh size).

#### Simulation of Sensors in Trial Insert

A third finite element model was created to simulate the interaction between the ball-like femoral head and socket i.e. the cup or acetabular component (Fig. S13), including the simulated forces at different points in the cup when a certain force is applied by the head. This model determines the theoretical force that each sensor should be reading, to compare to the actual response of the sensors in practice. Fig. S13a shows a schematic of the loading setup in the model – the ball-like femoral head applies a force $F$ to the cup-like acetabular part of the implant.

Fig. S13b shows how the pressure $P_0$ at a point on the internal surface of the cup varies with angle $\theta$ (in radians) and magnitude $F$ of the applied force. $P_0$ increases with applied load, and decreases linearly with angle $\theta$ from the force axis, resulting in the expression:

$$P_0 = P_{max}\left(1 - \frac{\theta^2}{\pi^2}\right) \quad (1)$$

where the pressure is at a maximum ($P_{max}$) at an angle of 0°. This expression agrees with the paraboloidal theory of contact between a sphere and a cup as established by Calonius and Saikko[27]. Fig. S13c shows a heat map of $P_0$ at $\theta$ = 0° and 30°. The model is used to carry out proof-of-concept simulations of the balance within the joint geometry. Fig. S13d shows the relative pressures exerted on the sensor reservoirs when a force is applied directly onto one of the reservoirs, at $\theta$ = 30°. This model will be used as a guide in the calibration of sensors in the hip implant geometry.

To determine the effect of incorporating sensors into the cup grooves, a 5 mm x 5 mm square section of the HDPE cup was replaced with PDMS (Fig. S13e) at a cup angle $\theta = 30°$, to represent a sensor embedded into the cup. The pressure $P_0$ was approximately 3.6 times smaller on the PDMS section ($P_0$ = 96 kPa) than on an HDPE section at the same cup angle ($P_0$ = 344 kPa), which represents a section of the cup made from the cup material with no embedded sensor. This indicates that the stiffer cup material is shielding the sensors from the applied force with the sensor-cup combination acting like a composite material, allowing the sensors to remain intact at higher applied forces. Fig. S13e also indicates that the shielding effect does not significantly change the forces in the areas of the cup surrounding the PDMS square. This analysis was also compatible with the observation that the maximum operating applied force was considerably higher in this cup geometry, with a sensitivity value of approximately 0.03 pF $N^{-1}$ – compared to 0.06 pF $N^{-1}$ when calibrating the sensors when not housed in the cup (see Fig. S2).

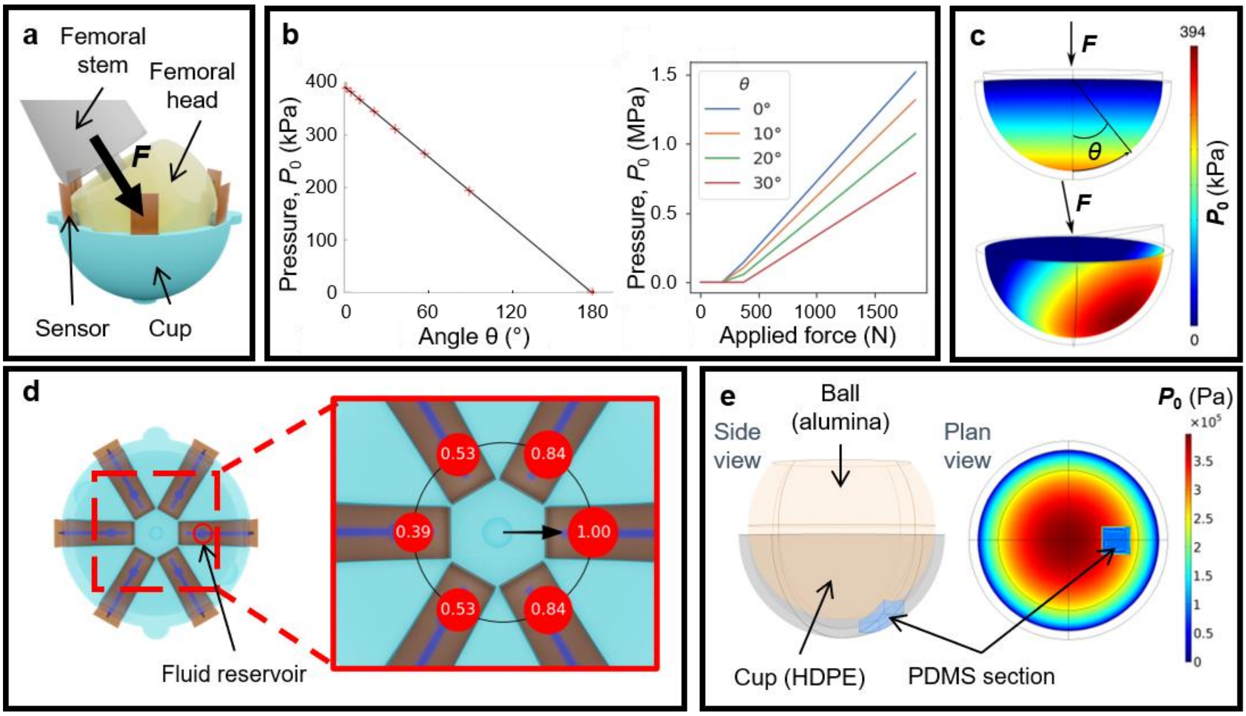


**Figure S13 | Modelling the loading of sensors incorporated into implant acetabular component / cup geometry.** **a** Schematic of angled loading; a force $F$ is applied normal to a sensor's fluid reservoir, at 30° to the vertical. **b,c** Finite Element Modelling simulations showing pressure $P_0$ in the cup as a function of applied force $F$ and the cup angle between $F$ and the cup normal, $\theta$. **d** Plan view of the cup, showing an array of six sensors and their fluid reservoirs. The force is applied normal to one reservoir, at $\theta$ = 30°, and the pressures shown are normalised by the pressure exerted on the reservoir closest to the contact point with the ball (femoral head). **e** Effect of replacing a 5 mm$^2$ section of the cup with PDMS, to show the effect of the cup shielding the sensors from the applied force.

#### *PDMS-Based Sensor Fabrication*

**Optimisation of printing method.** The group's previous work into fabricating the PDMS part of the device consisted of an AJP-based method, whereby NaCl salt was printed to form a sacrificial mold, PDMS was poured on top and cured, the resulting PDMS chip peeled off and the NaCl dissolved away[13]. This method reliably reproduced the square reservoir design, but the NaCl was difficult to dissolve as a PDMS-NaCl foam was deposited at the surface of the mold, often blocking the reservoir and making it difficult to fill with liquid. The foam can be removed, but this issue limits scalability and device reliability. In this paper, we present a new method. Molds were printed using the Formlabs Form 3 SLA printer and used to produce the devices as detailed in the Methods section. Although the SLA printer has a lower resolution (25 μm) compared to the AJP (10 ± 1 μm[14]), and the SLA molds took longer to produce (18 hours for a single mold for SLA, vs 3 hours for six molds for AJP), the SLA molds could be reused, which significantly decreased sensor production time overall. One disadvantage is that a new mold must be produced for each modification to the design. However, perhaps the greatest advantage of SLA for this application is that the microfluidic chip can be printed directly, using Flexible Resin for example, without the need of a mold.

### *Experimental Characterisation and Optimisation of PDMS-Based Sensor Design*

#### Sensor Calibration and Design Optimisation

Experimental calibration of PDMS-based sensors with different microfluidic chip designs are shown in Fig. S14.

**Channel dimensions:** Increased channel width or height leads to reduced sensitivity, as the larger cross sectional area of the channel would mean that the length of electrodes covered by liquid would be smaller for a certain applied force. Simulations were conducted with the same length of liquid covering the electrodes, but different channel width and height - as predicted, the capacitance values were near-identical (within 2%) for all channel widths above 0.5 mm (the electrode separation) and channel heights above 0.3 mm. Experimental results may also indicate this, as shown in Fig. S14. When comparing 0.25 mm- and 0.5 mm-wide channels, the former has a plateau of low sensitivity followed by the fluid reaching the end of the channel. Therefore, the channels could be made wider to incorporate a larger volume of liquid and go to higher forces, at the cost of sensitivity.

**Reservoir dimensions:** Increasing the width and height of the reservoirs leads to greater sensitivity, as more fluid is expelled at a certain force. However, taller reservoirs had a smaller rate of fluid expelled per unit force, meaning that they will be empty at higher forces. This relationship did not hold for wider reservoirs, as can be seen from a plateau in sensitivity at low forces. Increasing the reservoir width increases the difficulty in filling the reservoir completely – often the fluid enters the channel before filling the reservoir, leading to air bubbles.

**Reservoir shape:** In the group's previous work[13], a square reservoir was used. In this paper, it has been judged that circular cross-section reservoirs have the advantage of being easier to fill with fluid (preventing air bubbles) with the change in reservoir shape also having a insignificant effect on the sensitivity.

**Presence of Columns:** In the group's previous work[13], columns were used to provide mechanical stability and prevent collapse of the channel. For Flexible Resin-based devices the columns are no longer needed, perhaps in part due to the stress shielding provided by the implant geometry, and due to the increased stiffness of Flexible Resin. Furthermore, when fabricating the PDMS-based sensors it was difficult to fill in the reservoir without producing air bubbles in part due to the presence of the columns obstructing fluid flow down the sides of the reservoir. This issue causes inconsistent readings and lack of sensor consistency of measurements.

**Square vs Round columns:** This was investigated during modelling, but has not been verified experimentally yet. Modelling results are as follows: the increase in volume of the reservoir (i.e. the rounded columns are smaller than the square ones because their diameter = side of square) has more of an effect than the reservoir shape. Peak stress concentrations near reservoir columns reduced from 1.64 MPa to 981 kPa under 6 N of force.

The optimal design changes to make based on these results are to increase the channel and reservoir height. Increasing the length is also an option, but to achieve a large spatial resolution the sensor must be as small as possible. Options include creating a serpentine channel and electrodes.

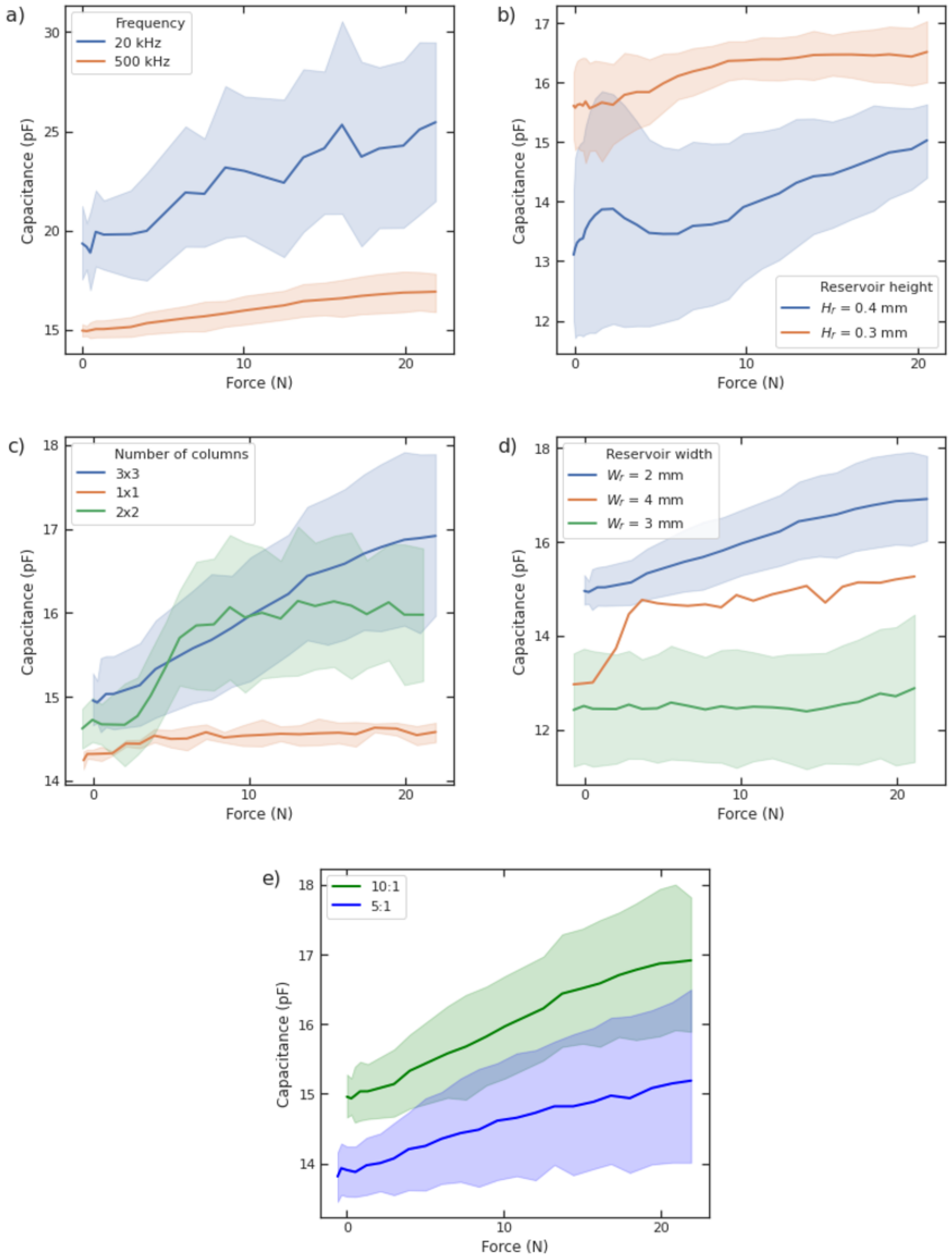


**Figure S14 | Experimental calibration of PDMS-based sensors up to 20 N, varying different sensor parameters.** The shaded area represent the 95% confidence interval from repeat experiments (repeat data was not available for some samples). **a** Frequency: A smaller measurement frequency results in a greater sensitivity but smaller signal to noise ratio. **b** Reservoir and channel height: a larger reservoir increases the sensitivity. **c** Number of columns: increasing the number of columns increases the sensitivity. **d** Reservoir width: increasing the reservoir width has little effect on the sensitivity. **e** Curing ratio: 10:1 and 5:1 curing ratios have similar sensitivities.

#### Fatigue Properties

The fatigue experiment results can be seen in Fig. S15 - the sensor lasted up to at least 100 cycles without significant change in device performance. Previously, it has been shown that these sensors are durable over at least 2000 cycles[13].

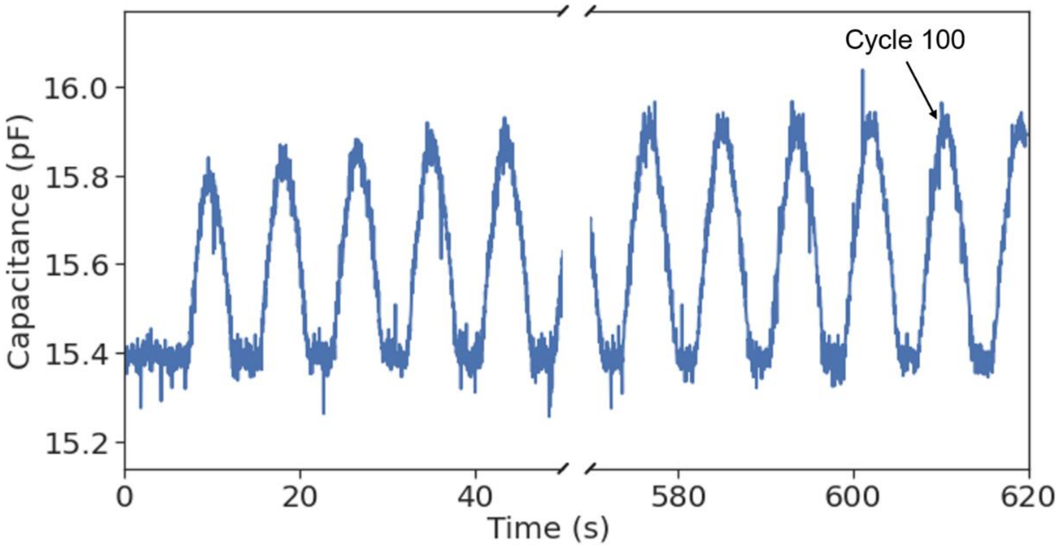


**Figure S15 | Effect of repeated loading on sensor performance.** The sensor lasted up to at least 100 cycles.

### Additional Mechanical Testing of Flexible Resin-Based Sensors in Trial Insert

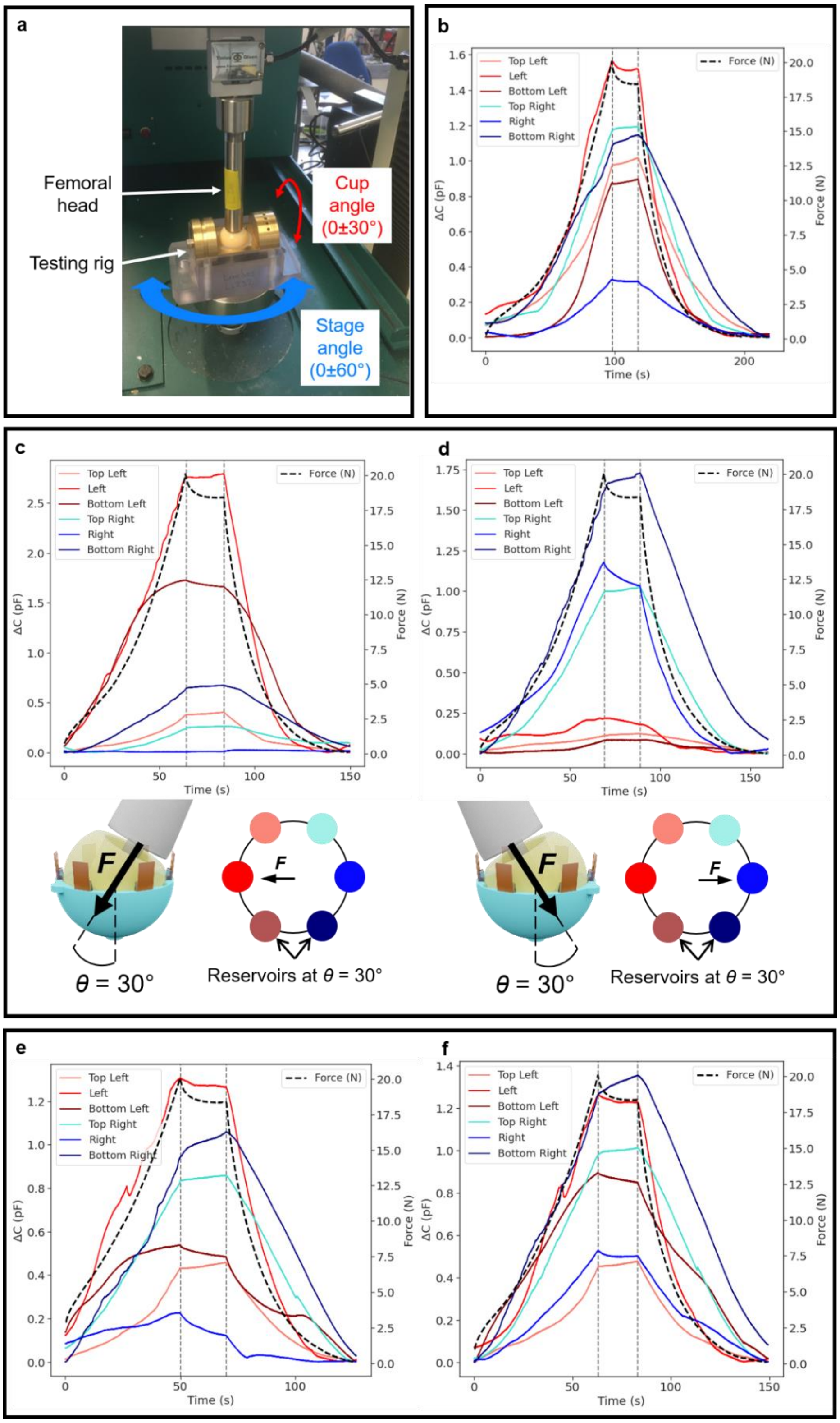


**Figure S16 | Effect of varying cup and stage angles. a** Experimental testing of the rig setup. The cup angle is the angle between the applied force and the normal to the plane of the cup. The stage angle is the lateral rotation of the rig. **b** Capacitance change with applied force (up to 20 N) over time for six sensors in the testing rig, with the cup and stage angles at 0°. The sensors are labelled based on their relative position in the cup. **c,d** Rotating the cup angle $\theta$ to ±30°. **e,f** Rotating the stage angle to ±60°.

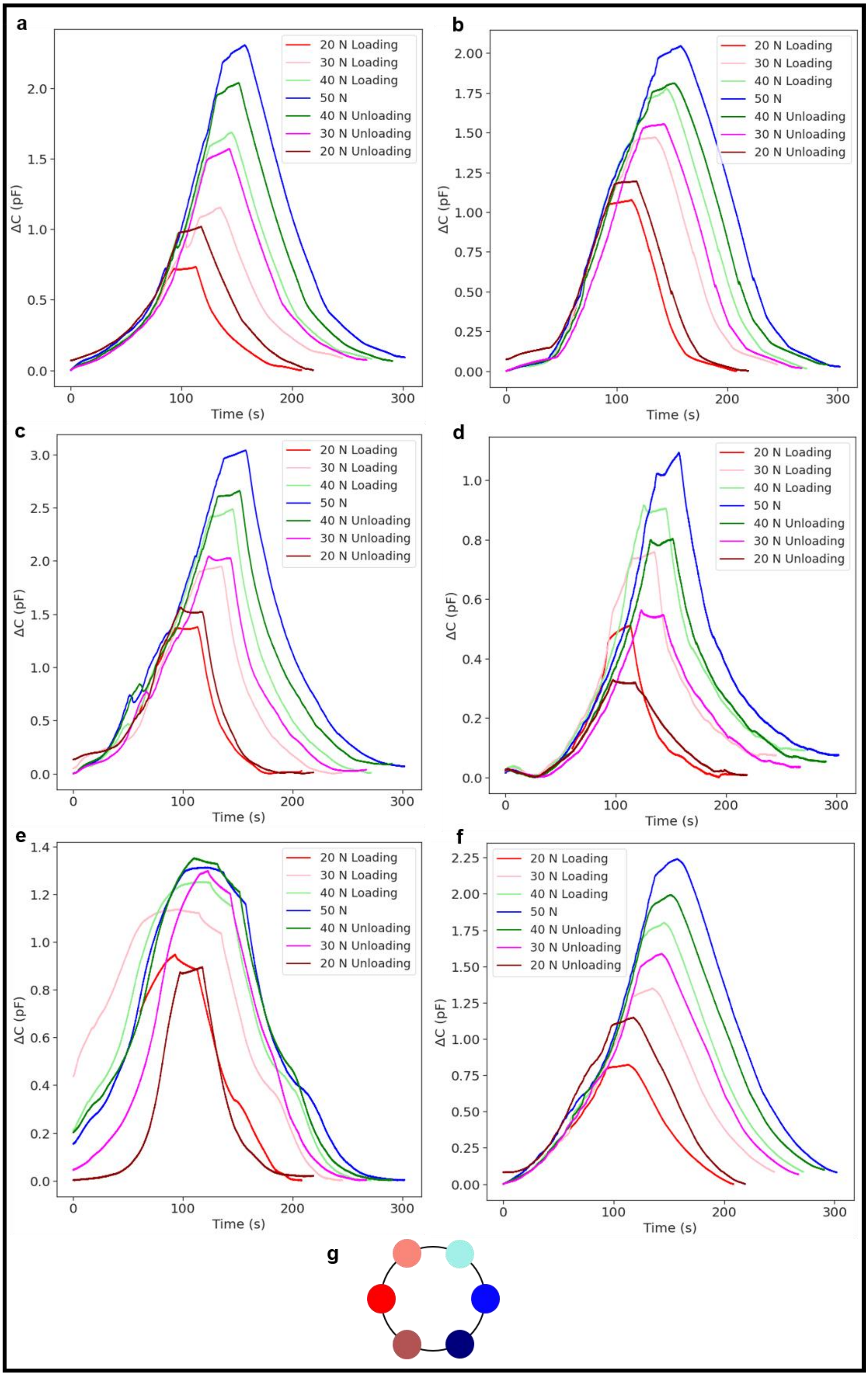


**Figure S17 | Repeated loading of six sensors,** in the **a** top left, **b** top right, **c** left, **d** right, **e** bottom left and **f** bottom right positions in the cup. The maximum force reached in a loading cycle was increased from 20 N to 50 N in steps of 10 N, then decreased to 20 N. **g** The sensors (represented by coloured dots) were located 60° apart from each other.
